# Pharmacological inhibition of PAI-1 alleviates cardiopulmonary pathologies induced by exposure to air pollutants PM_2.5_

**DOI:** 10.1101/2021.02.17.431681

**Authors:** Asish K Ghosh, Saul Soberanes, Elizabeth Lux, Meng Shang, Raul Piseaux-Aillon, Mesut Eren, G.R. Scott Budinger, Toshio Miyata, Douglas E Vaughan

## Abstract

**OBJECTIVE:** Exposure to air pollutants leads to the development of pulmonary and cardiovascular diseases, and thus air pollution is one of the major global threats to human health. Air pollutant particulate matter 2.5 (PM_2.5_)-induced cellular dysfunction impairs tissue homeostasis and causes vascular and cardiopulmonary damage. To test a hypothesis that elevated plasminogen activator inhibitor-1 (PAI-1) levels play a pivotal role in air pollutant-induced cardiopulmonary pathologies, we examined the efficacy of a drug-like novel inhibitor of PAI-1, TM5614, in treating PM_2.5_-induced vascular and cardiopulmonary pathologies.

**APPROACH AND RESULTS:** Results from biochemical, histological, and immunohistochemical studies revealed that PM_2.5_ increases the circulating levels of PAI-1 and thrombin and that TM5614 treatment completely abrogates these effects in plasma. PM_2.5_ significantly augments levels of pro-inflammatory cytokine IL-6 in bronchoalveolar lavage fluid, and this also can be reversed by TM5614, indicating its efficacy in amelioration of PM_2.5_-induced increases in inflammatory and pro-thrombotic factors. TM5614 reduces PM_2.5_-induced increased levels of inflammatory markers Mac3 and pSTAT3, adhesion molecule VCAM1, and apoptotic marker cleaved caspase 3. Longer exposure to PM_2.5_ induces pulmonary and cardiac thrombosis, but TM5614 significantly ameliorates PM_2.5_-induced vascular thrombosis. TM5614 also reduces PM_2.5_-induced increased blood pressure and heart weight. *In vitro* cell culture studies revealed that PM_2.5_ induces the levels of PAI-1, type I collagen, fibronectin, and SREBP-1/2, a transcription factor that mediates profibrogenic signaling, in cardiac fibroblasts. TM5614 abrogated that stimulation, indicating that it may block PM_2.5_-induced PAI-1 and profibrogenic signaling through suppression of SREBP-1. Furthermore, TM5614 blocked PM_2.5_-mediated suppression of Nrf2, a major antioxidant regulator in cardiac fibroblasts.

**CONCLUSIONS:** Pharmacological inhibition of PAI-1 with TM5614 is a promising therapeutic approach to control air pollutant PM_2.5_-induced cardiopulmonary and vascular pathologies.

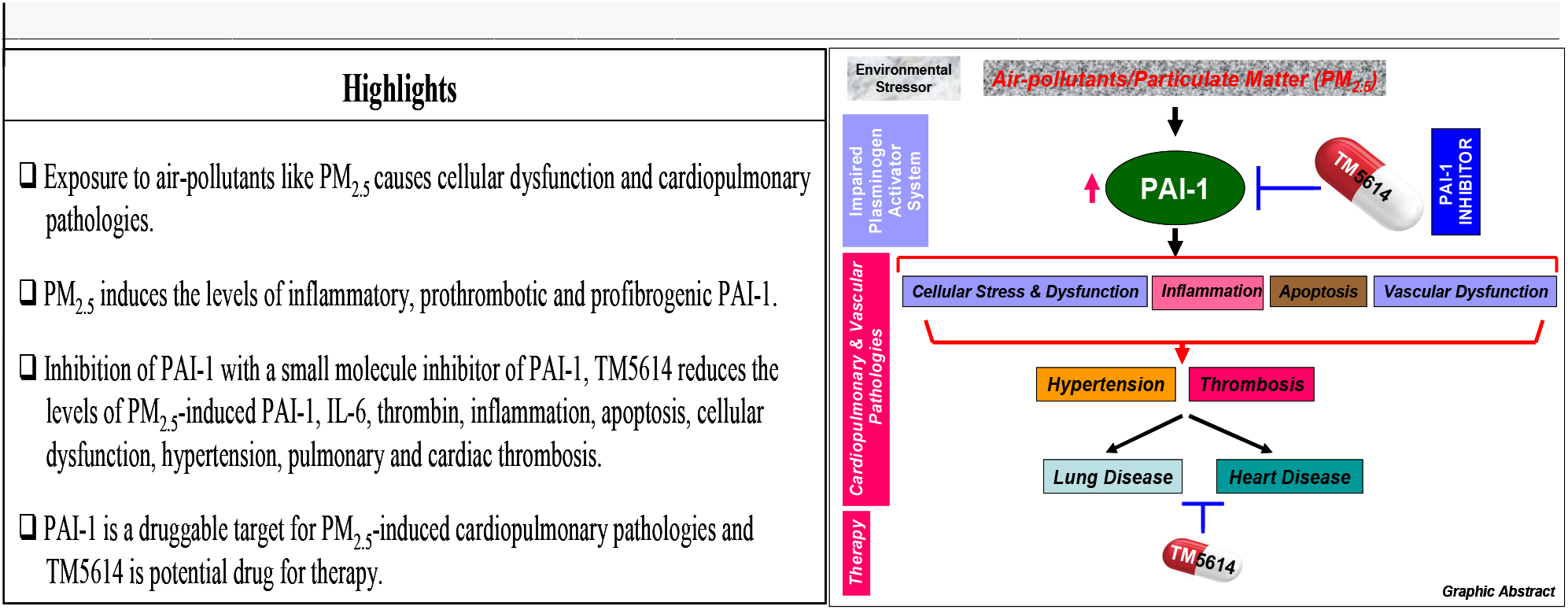

## INTRODUCTION

Air pollution-associated toxicity is a major threat to human health. Exposure to air pollutants leads to development of pulmonary and cardiovascular disease-related morbidity and mortality. Environmental pollutants like ozone, nitrogen oxide, and particulate matter (PM) induce oxidative stress, which leads to cardiopulmonary vascular diseases (CPVDs) including thrombosis, hypertension, arteriosclerosis, arrhythmias and, myocardial infarction—affecting millions of people worldwide.^1,2^ PM is the most dangerous environmental pollutant, as these solid particles contain a wide variety of toxic substances including sulfates, nitrates, hydrocarbons, heavy metals, and bacteria. Based on its aerodynamic diameter, PM has been classified as coarse (PM_10_), fine (PM_2.5_), and ultrafine (PM_0.1_).^3,4^ The detrimental effects of PM on the cardiopulmonary/vascular system stem from chronic or acute exposure-induced oxidative stress and systemic inflammation, cellular dysfunction, and impaired tissue homeostasis.^1–8^ While the acute effects of PM may be direct, involving rapid crossover from the lung epithelium into the circulation, the chronic effects of PM involve generation of pulmonary oxidative stress, systemic inflammation, and cellular dysfunction.^9^ With time, both direct and indirect effects of PM facilitate the onset and progression of CPVD pathologies. However, the exact molecular mechanism by which environmental pollutants like PM_2.5_ contribute to cardiopulmonary vascular diseases is not fully understood.

Plasminogen activator inhibitor-1 (PAI-1), a secretory protein that inhibits t-PA/uPA serine proteases, regulates the tissue fibrinolytic system, and its deregulation contributes to different diseases, including CPVDs like thrombosis and accelerated cellular senescence and aging.^10–14^ Here, we tested a hypothesis that elevation of PAI-1 plays a pivotal role in PM-induced CPVDs by inhibiting t-PA, increasing oxidative stress, and promoting cellular dysfunction and cardiopulmonary vascular injury; conversely, pharmacological inhibition of PAI-1 ameliorates PM-induced CPVDs. The strong rationale behind formulating this hypothesis comes from the previous reports that (i) the levels of PAI-1 are significantly elevated in oxidative stress-induced cardiac cells, (ii) elevated PAI-1 is a key determinant of premature cellular senescence and dysfunction in different pulmonary and cardiac cells, and (iii) pharmacological inhibition of PAI-1 effectively blocks oxidative stress-induced cellular dysfunction through suppression of key senescence mediators.^12–14^ In addition, the levels of PAI-1 are significantly elevated in bronchoalveolar lavage fluid (BALF) in mice, and in lung and heart tissues from rats, exposed to PM_2.5_ and are associated with inflammation and apoptosis.^15,16^ These findings strongly predict a therapeutically relevant function of PAI-1 inhibition in preventing PM_2.5_-induced CPVDs. In this study, we tested the efficacy of a novel druglike small molecule inhibitor of PAI-1, TM5614, in amelioration of air pollutant PM_2.5_-induced CPVD pathologies. The results of the present investigation reveal that pharmacological inhibition of PAI-1 in mice is associated with suppression of PM_2.5_-induced inflammatory and prothrombotic factors and thus of CPVD pathologies including vascular thrombosis.

## MATERIALS AND METHODS

All data and supporting materials have been provided with the published article and in the data supplement.

### Animal studies

Wild-type, PAI-1 heterozygous, and PAI-1 knockout C57Bl/6 mice (9-12 weeks old) were used for this study. All mouse protocols were approved by the Northwestern University Animal Care and Use Committee. Wild-type male mice were divided into 4 groups. Group 1: mice were fed with regular chow and received 50 μl of phosphate-buffered saline (PBS as control) (25 μl X2) by intratracheal instillation. Group 2: mice were fed with regular chow and received particulate matter 2.5 (PM_2.5_ in PBS) (the National Institutes of Standards and Technology, Gaithersburg, MD, USA; NIST SRM 1649) (50 μg-200μg/mouse; 50 μg or 100 μg or 200 μg in 50 μl PBS; 25 μl X2) by intratracheal instillation. Doses of PM_2.5_ were selected based on well accepted published literature.^15–21^ Group 3: mice were fed with chow containing TM5614 (10 mg/kg) for 6 days and then received PBS (25 μl X2 control) and TM5614 chow. Group 4: mice were fed with chow containing TM5614 (10 mg/kg) for 6 days and then received PM_2.5_ (50-200 μg/mouse in 50 μl PBS; 25 μl X2) and TM5614 chow. After 24 h or 72 h (short-term exposure to PM_2.5_ :50 and 200 μg/mouse/week or PBS in the presence or absence of TM5614) or 4 weeks (longterm exposure to PM_2.5_: 50-200 μg/mouse/week or PBS in the presence or absence of TM5614), mice were sacrificed and plasma and BALF (short-term exposure only) were collected and processed for biochemical analysis. For PAI-1 genetic deficiency study, batches of wild-type, PAI-1 heterozygous, and PAI-1 knockout male and female mice were fed with regular chow and treated either with PM_2.5_ (50 μg/mouse/week for 4 weeks) or PBS as controls. Hearts and lungs from short-term exposed mice (72 h to study the early pathological events) and long-term-exposed mice (4 weeks to study late pathological events) were collected and used for histological and immunohistochemical analysis.

### Measurement of PAI-1 and thrombin-antithrombin (TAT) complex levels in plasma

Plasma was collected from the 4 groups of wildtype mice, and 50 μl of TAT standard and each sample were used in duplicate for TAT assays using a mouse ELISA kit for TAT (cat no ab137994, Abcam). Similarly, PAI-1 standard and each plasma sample were used in duplicate for PAI-1 assays using a mouse PAI-1 ELISA kit (cat no ab197752, Abcam).

### Measurement of IL-6 in BALF

BALF was collected from each group of mice after 24 h of treatment as indicated above, and 50 μl of IL-6 standard and each sample were used in duplicate for IL-6 protein measurement using a Quantikine IL-6 mouse ELISA kit (cat no M6000B, R&D).

### Blood pressure and echocardiography

To monitor the effect of PM_2.5_ exposure on blood pressure, both systolic and diastolic blood pressure were measured in conscious mice using a non-invasive tail-cuff system (CODA Non-Invasive Blood Pressure, Kent Scientific Corp.). Transthoracic two-dimensional M-mode echocardiography was performed using a Vevo 3100 device (VisualSonics, Toronto, Canada) equipped with a MX550D transducer. Echocardiographic studies were performed after 4 weeks of PM_2.5_ exposure following manufacturer’s instruction. M-mode tracings were used to measure left ventricular (LV) wall thickness, LV dimensions, LV volume, and LV mass. Percent fractional shortening (% FS) and ejection fraction (% EF) were also determined. The mean value of at least 3-5 cardiac cycles was used to determine the measurements for each animal in all four groups. Post-mortem heart weight and body weight of each mouse in all four groups were recorded and compared.

### Histology

Heart and lung tissues were collected and fixed in 10% formalin overnight at room temperature, followed by fixation in 70% ethanol at 4°C overnight, and then processed for embedding in paraffin. Paraffin-embedded heart and lung tissues were subjected to microtome sectioning and processed for histological analysis. For morphology studies, the tissues were processed for hematoxylin and eosin (H&E) staining. The H&E staining of specimens was performed on a fully automated platform (Leica Autostainer XL) using Harris hematoxylin (Fisherbrand cat no.245-651) and eosin (Leica cat no. 3801600). The levels of collagen deposition and thrombotic blood vessels in lung and heart tissues were determined by H&E and Masson trichrome staining. Photographs were taken with an Olympus DP71 camera. The levels of matrix deposition were estimated by Fiji-ImageJ software.

### Immunohistochemistry

All the immunohistochemistry staining was completed using chromogenic enzyme substrate reactions with DAB (DAKO,cat no.K3468). Formalin-fixed paraffin-embedded tissue sections (4 μm) were first deparaffinized on an automated platform (Leica Autostainer XL). After deparaffinization, slides were subjected to an antigen retrieval step using a sodium citrate solution at pH 6 in a pressure cooker (Biocare). Primary antibody was then added manually and incubated at 4 °C overnight in a humid chamber. Secondary antibody incubation and chromogenic reactions were then performed using an automated system (Biocare Intellipath). Once the staining was complete, specimens were counterstained with hematoxylin. Slides were then coverslipped using a xylene-based mounting medium (Leica Micromount). The antibodies used in this study are as follows: Mac3: BD Biosciences cat no.553322; concentration 1:5000. Antigen retrieval: pH 6 sodium citrate, 11°C for 10 min. pSTAT3: Cell Signaling, cat.no.9145; concentration 1:250. Antigen retrieval: pH 6 sodium citrate, 11 °C for 10 min. Cleaved Caspase 3: Cell Signaling cat .no.9661; Concentration 1:500; Antigen Retrieval: Ph6 Sodium Citrate, 11°C for 10min.VCAM1: Abcam ab134047, concentration 1:500. Antigen retrieval: pH 6 sodium citrate, 11 °C for 20 min. Photographs were taken using a light microscope. The sum of several fields per heart and lung tissue from each mouse were used for staining intensity measurement using ImageJ software, and the levels of statistical significance were determined.

### Cell cultures and treatment with PM_2.5_ and TM5614

Primary cultures of human cardiac fibroblasts were grown in Dulbecco’s Modified Eagle Medium (DMEM) containing 10% fetal bovine serum (FBS) and 1% penicillin and streptomycin. To test the efficacy of PAI-1 inhibitor TM5614 in suppression of PM_2.5_-induced cardiac fibroblast dysfunction, confluent cultures of human cardiac fibroblasts were subcultured in 12-well clusters. After 24 h, media were replaced with 0.1% FBS-containing DMEM for 3 h. Cells were pretreated with TM5614 (10 μM) or DMSO (vehicle control) for 2 h followed by treatment with PM_2.5_ (50 μg/ml) in triplicate for another 2 h. For the other set of experiments, cells were pretreated with TM5614 (10 μM) or DMSO in triplicate for 24 h. After 24 h, media were replaced with 0.1% FBS-containing DMEM and treated with TM5614 (10 μM) or DMSO and PM_2.5_ (50 μg/ml) in triplicate for another 24 h.

### Preparation of cell lysates and western blot analysis

At the end of treatment with PM_2.5_ in the absence and presence of PAI-1 inhibitor TM5614, supernatants from control and treated wells were collected. The cell lysates were prepared using RIPA lysis buffer containing protease inhibitor cocktail (cOmplete) and phosphatase inhibitor cocktail (PhosSTOP) (Sigma). Cell lysates were pooled from 3 wells for each control and treatment group and equal amounts of protein were subjected to gel electrophoresis, transferred to PVDF membranes, and subjected to western blotting using antibodies against PAI-1 (Molecular Innovations, Inc.), type I collagen (Southern Biotech), fibronectin, SREBP1, SREBP2, Nrf2 (Abcam), and α-tubulin (GenScript), with HRP-tagged corresponding secondary antibodies. The membranes were developed with enhanced chemiluminescence reagents (Luminata Forte, Millipore), and images of protein bands were captured using a BIO-RAD ChemiDoc XRS system (BIO-RAD, CA).

### Statistical analysis

Data are presented as mean ± SEM. The significance of differences between controls and experimental groups was determined by statistical analysis using Student’s t-test, where a value of P<0.05 was considered statistically significant. Statistical analyses were performed with GraphPad Prism (GraphPad Software Inc., San Diego, CA).

## RESULTS

### PAI-1 inhibitor TM5614 reduces PM_2.5_ exposure-induced plasma levels of PAI-1 and thrombin in mice

Four groups of mice described in the Methods section were used in this study: group 1, controls; group 2, PM_2.5_ only; group 3, TM5614 only; and group 4, TM5614 and PM_2.5_. After 24 h or 72 h, mice were sacrificed, and plasma was used for PAI-1 and TAT assays. The results revealed that exposure to PM_2.5_ (50 μg/mouse/once) for 24 h increases the levels of PAI-1 significantly, and TAT complex insignificantly in plasma. Importantly, pretreatment of mice with PAI-1 inhibitor TM5614 completely abrogates the PM_2.5_-induced increased levels of PAI-1 and TAT in plasma (Figure 1A, B). Similarly, exposure to PM_2.5_ (200 μg/mouse/once) for 72 h significantly stimulates the levels of TAT in plasma, and TM5614 abrogates that stimulation (Figure 1C). These results indicate that PAI-1 inhibitor TM5614 is highly efficient in reduction of PM_2.5_-induced PAI-1 and thrombin, which contribute to vascular pathologies like thrombosis.

**Figure 1.**
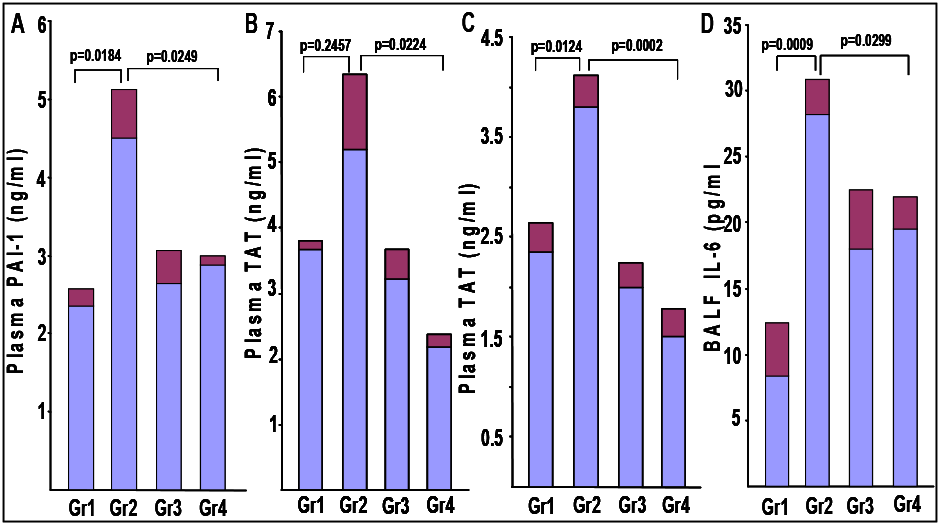
PAI-1 inhibitor TM5614 inhibits the levels of PM_2.5_-induced PAI-1 and TAT in plasma and IL-6 in BALF. Plasma collected from 4 groups of mice (n=4-8) was used in duplicate for PAI-1 and TAT assays. Day 1-6: TM5614 (10 mg/kg); Day 7: PM_2.5_ (50 μg/mouse) or PBS by intratracheal instillation; Day 8: Plasma collected and processed for PAI-1 (**A**) or TAT (**B**) assay using an ELISA kit. (**C**) TAT levels in plasma from mice after 72 h treatment with PM_2.5_ (200 μg/mouse) and TM5614 (10 mg/kg) (n=4-7). (**D**) IL-6 levels in BALF from mice after 24 h treatment with PM_2.5_ (50 μg/mouse) and TM5614 (10 mg/kg) were determined in duplicate using an ELISA kit (n=7-8). Gr 1: PBS; Gr2: PM2.5; Gr3: TM5614; Gr4: TM5614+PM2.5.

### PAI-1 inhibitor TM5614 significantly inhibits PM_2.5_-induced IL-6 in BALF

BALF collected from the four groups of mice was used for determination of IL-6 levels as described in the Methods. Administration of PM_2.5_ (50 μg/mouse/ once) for 24 h significantly increases the levels of pro-inflammatory cytokine IL-6 in BALF and most importantly, PM_2.5_ fails to stimulate the levels of IL-6 in mice fed with TM5614, indicating the efficacy of PAI-1 inhibitor TM5614 in amelioration of PM_2.5_-induced lung inflammation (Figure 1D). The levels of IL-6 in BALF from TM5614-treated (Group 3) and PBS control (Group 1) mice are not significantly increased (p=0.1301).

### PAI-1 inhibitor TM5614 alleviates PM_2.5_-induced lung inflammation, apoptosis, and vascular pathologies

Wild-type mice were exposed to a single dose of PM_2.5_ (200 μg/mouse). After 72 h, lung tissues were collected and processed for morphological analysis by H&E staining and immunohistochemical analysis using antibodies against inflammatory markers Mac3 and pSTAT3, apoptotic marker cleaved caspase 3, and adhesion molecule VCAM1. Morphologically, PM_2.5-_exposed lungs (4 out of 7) are relatively denser compared to PBS controls (1 out of 4) and importantly, TM5614 partially reverses the PM_2.5_-induced lung morphological changes (Supplemental Figure s1, representative images are presented). Immunohistochemistry results revealed the presence of low levels of inflammation in PBS treated lungs. However, administration of PM_2.5_ increases the staining intensity of pSTAT3 and Mac3 compared to controls and mice treated with TM5614 alone. Treatment with PAI-1 inhibitor TM5614 modestly reduces PM_2.5_-induced pSTAT3 without reaching statistical significance (Figure 2A, representative images are presented and quantitative data are shown in Figure 2B) and significantly reduces the Mac3 (Figure 3A, representative images are presented and quantitative data are shown in Figure 3B) levels in lung tissues. These results are consistent with our observation that PM_2.5_ induces the level of pro-inflammatory cytokine IL-6 in BALF and TM5614 completely abrogates the PM_2.5_-induced increased levels of IL-6 in lungs.

**Figure 2.**
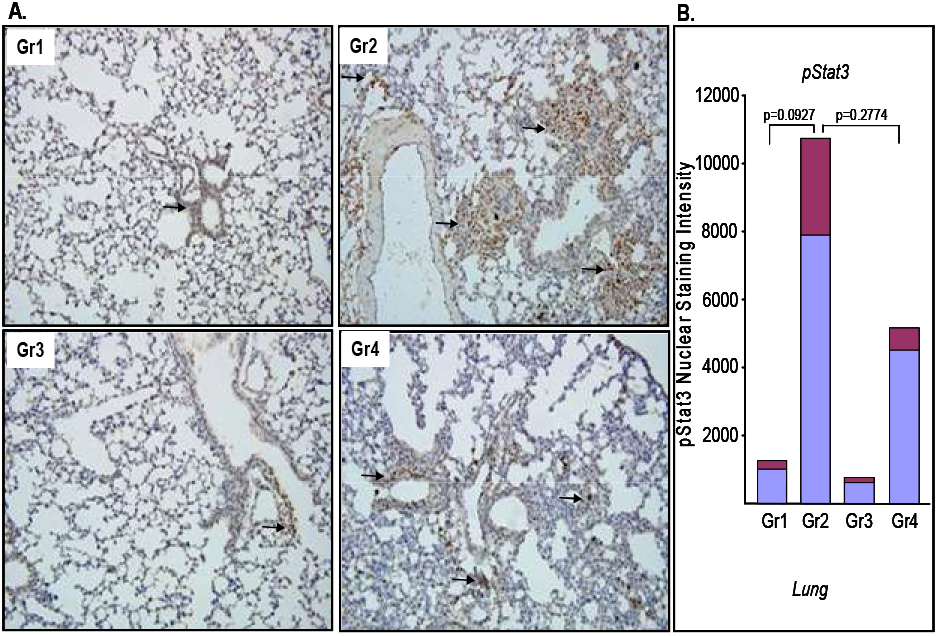
Effect of PAI-1 inhibitor TM5614 on PM_2.5_-induced inflammatory factor pSTAT3 in lungs. Lungs collected from 4 groups of mice (n=4-6) were processed for immunohistochemistry using an anti-pSTAT3 antibody. Day 1-6: TM5614 (10 mg/kg); Day 7: PM2.5 (200 μg/mouse) or PBS instillation; Day 10: Lungs were collected and processed for immunohistochemistry. Representative images are shown in (**A**). Images are reduced to 15% of original 20X images. The levels of nuclear pSTAT3 in several fields of each lung section were determined by ImageJ followed by statistical analysis. Quantitative data are shown in (**B**). Gr1: PBS; Gr2: PM_2.5_; Gr3: TM5614; Gr4: TM5614+ PM_2.5_.

**Figure 3.**
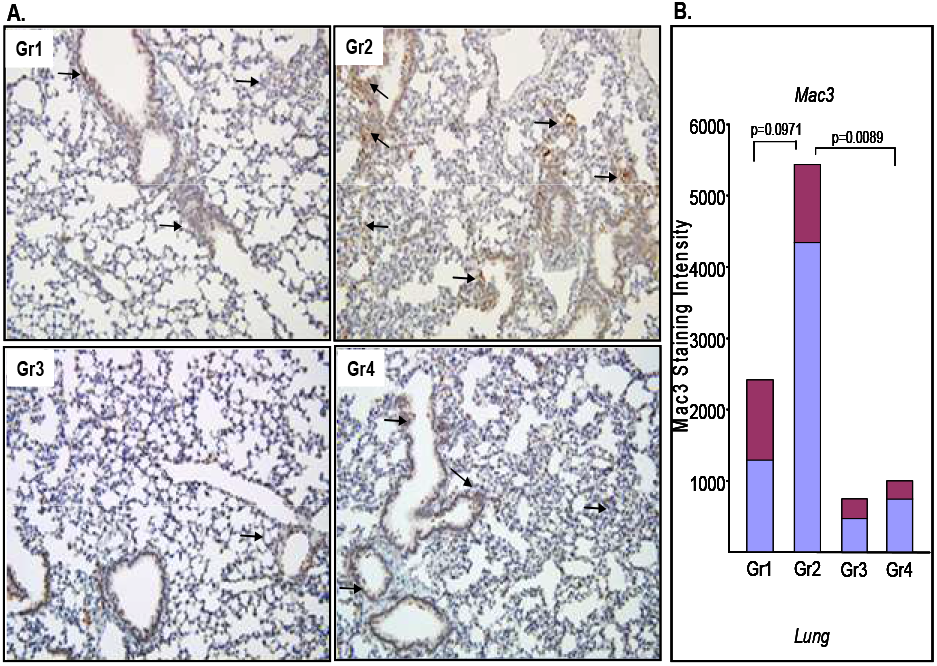
PAI-1 inhibitor TM5614 reduces PM_2.5_-induced inflammation in lungs. Lungs collected from 4 groups of mice (n=4-6) were processed for immunohistochemistry using an anti-Mac3 antibody. Day 1-6: TM5614 (10 mg/kg); Day 7: PM_2 5_ (200 μg/mouse) or PBS instillation. After 72 h, lungs were collected and processed for immunohistochemistry. Representative images are shown in (**A**). Images are reduced to 15% of original 20X images. The levels of Mac3 intensity in several fields of each lung section were determined by ImageJ software. Quantitative data are presented in (**B**). Gr1: PBS; Gr2: PM_2 5_; Gr3: TM5614; Gr4: TM5614+PM_2.5_.

Exposure to PM_2.5_ for 72 h significantly increases the levels of cleaved caspase 3 positivity in lungs (Figure 4A, representative image from each group; quantitative data are shown in Figure 4B). Importantly, TM5614 treatment reduces the levels of cleaved caspase 3 positivity in lungs. Next, we determined the levels of adhesion molecule VCAM1, known to be involved in inflammation and vascular dysfunction, in PM_2.5_-exposed murine lungs. Exposure to PM_2.5_ induces the levels of VCAM1, consistent with the previous observation that PM_2.5_-induced stress increases VCAM1 in endothelial cells.^22,23^ Most importantly, TM5614 prevents PM_2.5_-induced induction of VCAM1 (Figure 5A, representative images are presented from each group; quantitative data are shown in Figure 5B). Collectively, these results indicate that pharmacological inhibition of PAI-1 is an ideal approach to control air pollutant-induced inflammation and cellular apoptosis in murine lungs and that small molecule TM5614 effectively prevents air pollutant-induced lung pathological events.

**Figure 4.**
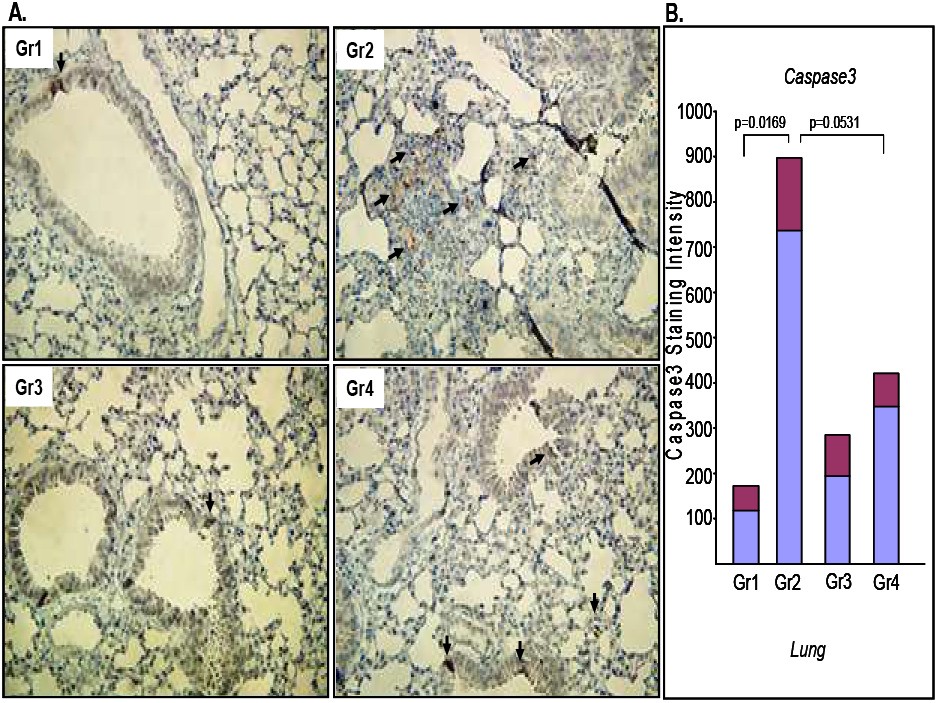
Effect of TM5614 on PM_2 5_-induced cellular apoptosis in lungs. Lungs collected from 4 groups of mice (n=4-6) were processed for immunohistochemistry using an anti-cleaved caspase 3 antibody. Mice were fed with TM5614 (10 mg/kg) for 6 days and on day 7, PM2.5 (200 μg/mouse) instillation or PBS was used as control. After 72 h, lungs were processed for immunohistochemistry. Representative images are reduced to 15% of original 40X images (**A**). The levels of cleaved caspase 3 in several fields of each lung were determined by ImageJ software. Quantitative data are shown in (**B**). Gr1: PBS; Gr2: PM_2.5_; Gr3: TM5614; Gr4: TM5614+PM_2.5_.

**Figure 5.**
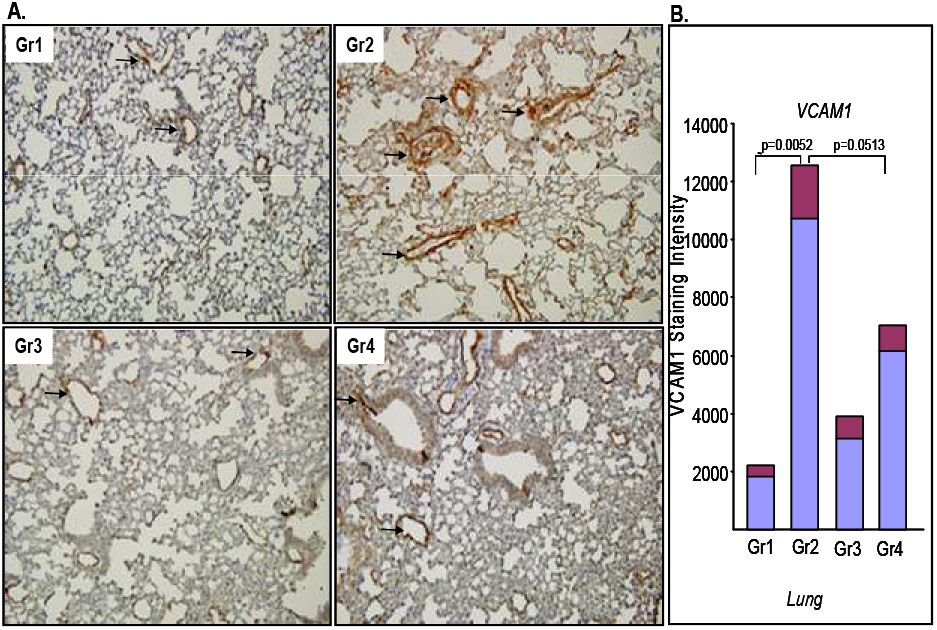
TM5614 reduces PM_2.5_-induced adhesion molecule VCAM1 in lungs. Lungs collected from 4 groups of mice (n=4-6) were processed for immunohistochemistry using an anti-VCAM1 antibody. Day 1-6: TM5614 (10 mg/kg); Day 7: PM_2.5_ (200 μg/mouse) or PBS instillation; Day 10: Lungs were processed for immunohistochemistry. Representative images are reduced to 15% of original 20X images (**A**). The levels of VCAM1 in several fields of each lung were determined by ImageJ. Quantitative data are shown (**B**). Gr1: PBS; Gr2: PM_2.5_; Gr3: TM5614; Gr4: TM5614+PM_2.5_.

### Effect of PAI-1 inhibitor TM5614 on PM_2.5_-induced cardiac inflammation and apoptosis

Hearts collected from the four groups of mice were processed for H&E staining and immunohistochemical analysis using antibodies against pSTAT3, Mac3, cleaved caspase 3, and VCAM1. There were no morphological changes in PM_2.5_-exposed hearts (Supplemental Figure s2). Immunohistochemistry results revealed that exposure to PM_2.5_ (200 μg/mouse/once) for 72 h increases the levels of cleaved caspase 3 in myocardial tissues and that TM5614 treatment reduces the levels of cleaved caspase 3 without attaining statistical significance (Figure 6A, representative image from each group; quantitative data are shown in Figure 6B). The levels of PM_2.5_-induced apoptotic cells in hearts are lower compared to lungs. While administration of PM_2.5_ significantly increases cardiac inflammation as evidenced by increased Mac3 staining, TM5614 modestly reduces inflammation (Figure 7A, representative image from each group; quantitative data are shown in Figure 7B). There is no difference in the levels of pSTAT3 in cardiac tissues derived from control and treated mice (Supplemental Figure s3A, representative image of each group, and quantitative data are shown in Figure s3B). While exposure to PM_2.5_ modestly stimulates the levels of VCAM1 in myocardial tissues, in the TM5614-treated mice, PM_2.5_ fails to stimulate adhesion molecule VCAM1 (Figure 8A, representative image from each group; quantitative data are shown in Figure 8B). These results signify the efficacy of PAI-1 inhibitor TM5614 in suppression of air pollutant-induced cardiovascular pathologies.

**Figure 6.**
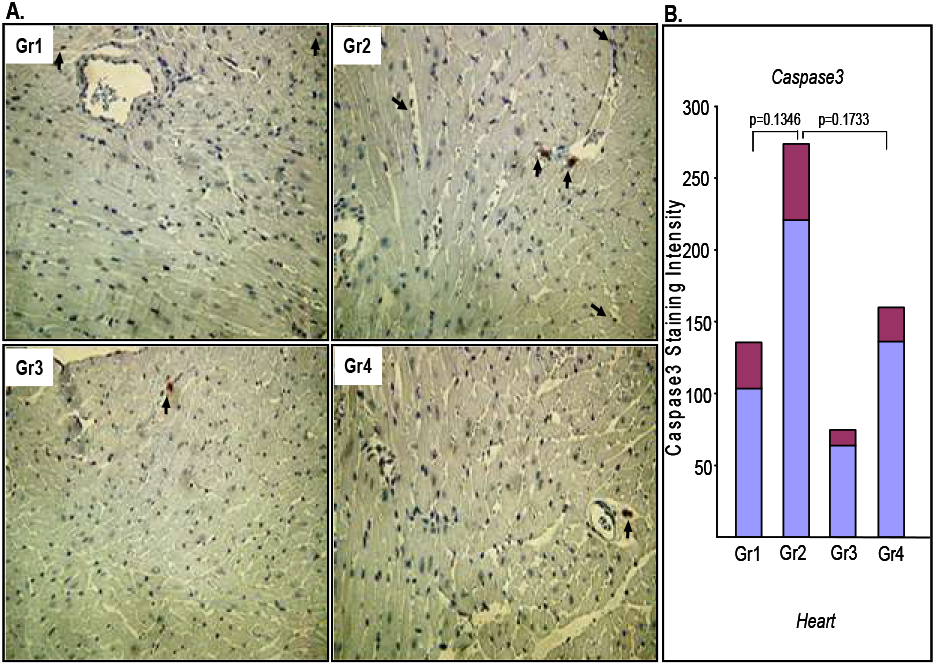
Effect of PAI-1 inhibitor TM5614 on PM_2.5_-induced cellular apoptosis in hearts. Hearts collected from 4 groups of mice (n=4-6) were processed for immunohistochemistry using an anti-cleaved caspase 3 antibody. Day 1-6: TM5614 (10 mg/kg); Day 7: PM_2.5_ (200 µg/mouse) or PBS instillation. After 72 h, hearts were collected and processed for immunohistochemistry. Representative images are reduced to 15% of original 40X images (**A**). The levels of cleaved caspase 3 in several fields of each heart section were determined by ImageJ. Quantitatve data are shown in (**B**). Gr1: PBS; Gr2: PM_2 5_; Gr3: TM5614; Gr4: TM5614+PM_2.5_.

**Figure 7.**
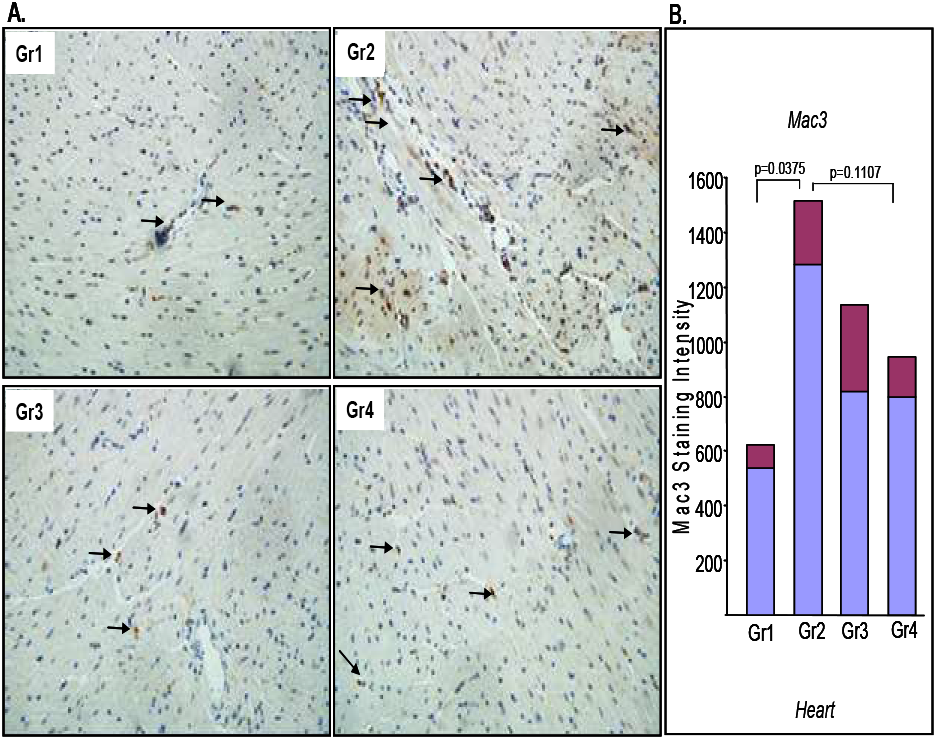
PAI-1 inhibitor TM5614 reduces PM_2.5_-induced inflammation in murine hearts. Hearts were collected from 4 groups of mice (n=4-6) and processed for immunohistochemistry using an anti-Mac3 antibody. Day 1-6: TM5614 (10 mg/kg); Day 7: PM_2 5_ (200 μg/mouse) or PBS instillation. After 72 h, hearts were processed for immunohistochemistry. Representative images are shown in (**A**). Images are reduced to 15% of original 40X images. The levels of Mac3 intensity in several fields of each heart were used for quantification using ImageJ. Gr1 : PBS; Gr2: PM_2.5_; Gr3 : TM5614; Gr4: TM56 1 4+PM_2.5_.

**Figure 8.**
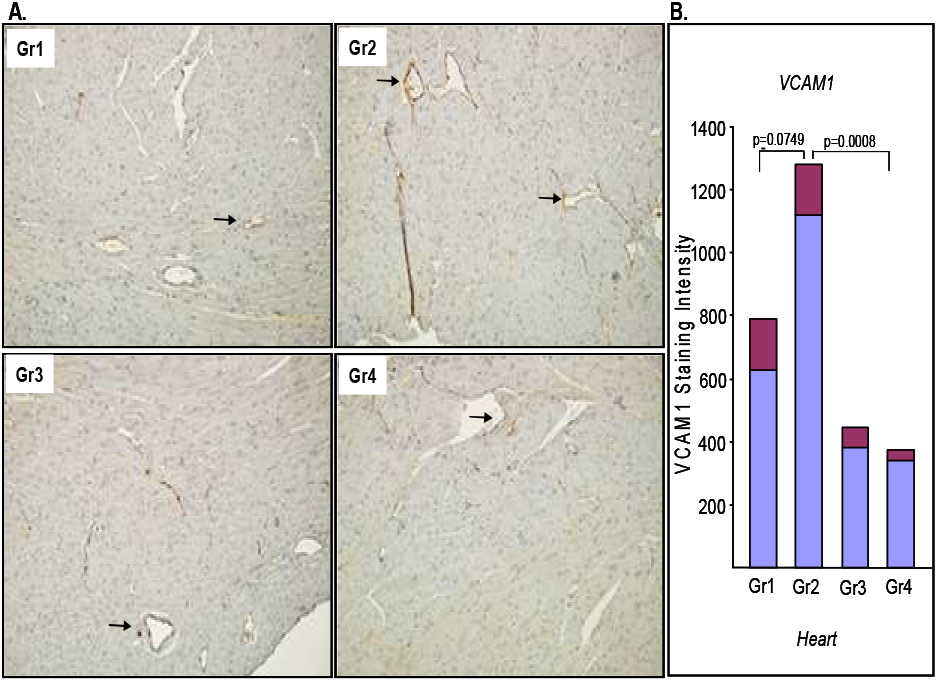
TM5614 reduces PM_2.5_-induced adhesion molecule VCAM1 in hearts. Hearts collected from 4 groups of mice (n=4-6) as indicated were processed for immunohistochemistry using an anti-VCAM1 antibody. Day 1-6: TM5614 (10 mg/kg); Day 7: PM_2.5_ (200 μg/mouse) or PBS instillation; Day 10: Heart sections were processed for immunohistochemistry.In (**A**), representative images are shown. Images are reduced to 15% of original 20X images. The levels of VCAM1 in several fields of each heart were determined by ImageJ software. Quantitative data are shown in (**B**). Gr1 : PBS; Gr2: PM_2.5_; Gr3: TM5614; Gr4: TM5614+PM_2.5_.

### Effect of long-term exposure to PM_2.5_ on blood pressure, cardiac hypertrophy, and cardiac matrix remodeling

Mice were divided into 4 groups. In Group1, mice were fed with regular chow and received PBS. Group 2, mice were fed with regular chow and received PM_2.5_ (100 μg/mouse/once per week for 4 weeks). Group 3, mice were fed with chow containing TM5614 (10 mg/kg) for 6 days and then received PBS for 4 weeks. Group 4, mice were fed with chow containing TM5614 (10 mg/kg) for 6 days and then received PM_2.5_ (100 μg/mouse/once per week for 4 weeks). At the end of the 4 weeks, blood pressure was determined, and each mouse was subjected to echocardiography. The results revealed that 4 weeks of PM_2.5_ exposure increases systolic pressure non-significantly and diastolic blood pressure significantly. Importantly, treatment of mice with PAI-1 inhibitor TM5614 reduces the PM_2.5_-induced blood pressure increase (Supplemental Figure s4A,B). Analysis of echocardiography data revealed that there is no significant differences in EF, FS, left ventricular anterior or posterior wall thickness-systolic or diastolic (LVAWs, LVAWd, LVPWs and LVPWd) within control and treated groups (data not shown). Post-mortem heart weight analysis reveals that PM_2.5_ (100 μg/mouse once per week for 4 weeks) induces cardiac hypertrophy based on increased heart weight/body weight ratio and that TM5614 decreases the PM_2.5_-induced cardiac hypertrophy (Supplemental Figure s4C). Collectively, these results suggest that TM5614 is effective in amelioration of PM_2.5_-induced cardiac pathologies including hypertension and cardiac hypertrophy. However, PM_2.5_ fails to stimulate fibrogenesis in heart tissue after 4 weeks of treatment (100 μg once per week for 4 weeks) (*data not shown*).

### PAI-1 inhibitor TM5614 reduces long-term PM_2.5_ exposure-induced pulmonary and cardiac thrombosis in mice

Mice were divided into 4 groups and treated for 4 weeks as described above. At the end of treatment, mice were sacrificed, and heart and lungs were collected. Lung and heart tissues were processed for H&E staining and Masson’s trichrome staining. Exposure to PM_2.5_ for 4 weeks significantly induces pulmonary thrombogenesis in mice (incidence of thrombus formation: 4 out of 7) compared to PBS controls (incidence of thrombus formation: 0 out of 4) and mice treated with TM5614 alone (incidence of thrombus formation: 0 out of 5). Most importantly, PM_2.5_ failed to induce pulmonary thrombosis in mice pretreated with PAI-1 inhibitor TM5614 (incidence of thrombus formation: 0/4) (Figure 9, representative image of each group). This result is highly significant, as it clearly indicates the beneficial effect of PAI-1 inhibitor TM5614 in improving vascular thrombosis, a major pulmonary pathology, in mice exposed to air pollutant PM_2.5_. However, PM_2.5_ fails to stimulate the pulmonary fibrogenesis significantly after 4 weeks of treatment (Supplemental Figure s5).

**Figure 9.**
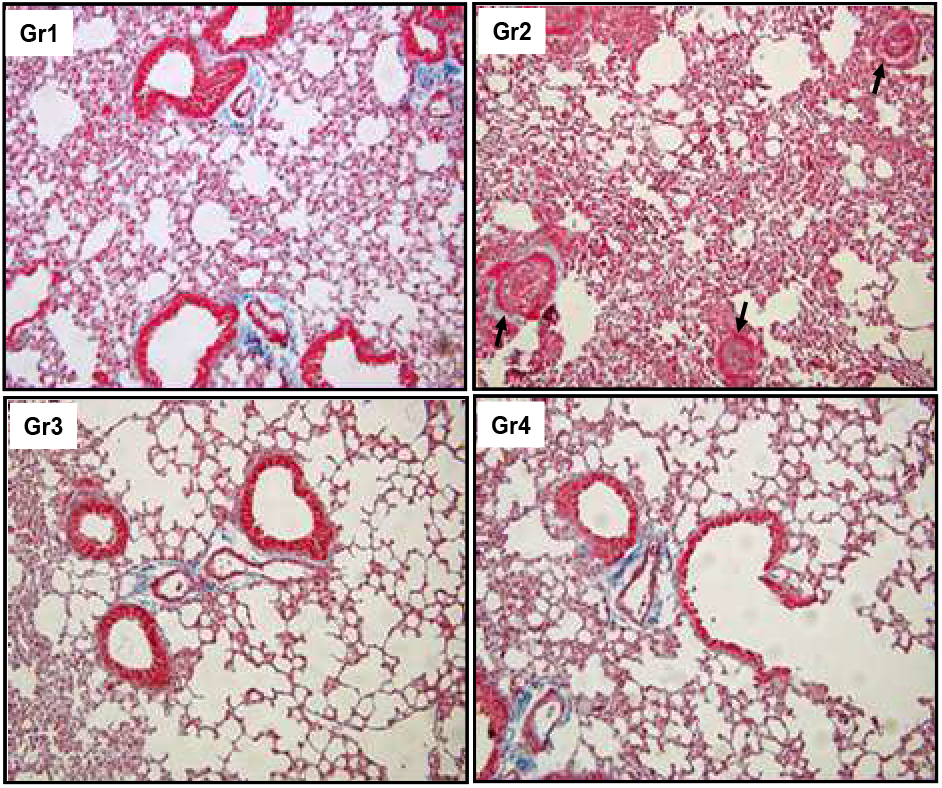
PM_2.5_ induces pulmonary thrombosis and PAI-1 inhibitor TM5614 prevents lung thrombogenesis. Mice were divided into 4 groups (n=4-7). In Group1, mice were fed with regular chow and received PBS. In Group 2, mice were fed with. regular chow and received PM_2.5_ (100 μg/mouse/once per week for 4 weeks). In Group 3, mice were fed with. chow containing TM5614 (10 mg/kg) for 6 days and then received PBS. In Group 4, mice were fed with chow containing TM5614 (10 mg/kg) for 6 days and then received PM_2.5_ (100 μg/mouse/once per week for 4 weeks). Representative image from each group are shown. Images reduced to 15% of original 20X images. An arrow indicates the presence of a thrombus.

Analysis of hearts from each group revealed that longer exposure to PM_2.5_ modestly induces cardiac thrombosis in mice, to a lesser extent than in lungs. However, PM_2.5_ failed to induce cardiac thrombosis in mice pretreated with PAI-1 inhibitor TM5614 (Supplemental Figure s6). These results are consistent with our observation that PM_2.5_ induces expression of PAI-1 and thrombin, the major contributors to thrombogenesis, and that TM5614 completely abrogates the PM_2.5_-induced increased levels of PAI-1 and thrombin in plasma (Figure 1). Additionally, these results further indicate that PAI-1 is an ideal druggable target for the treatment of air pollutant-induced cardio-pulmonary pathologies, including vascular thrombosis.

### PAI-1 heterozygous mice are partly protected from PM_2.5_-induced pulmonary thrombosis

As a complementary study to pharmacological inhibition of PAI-1 and its beneficial effect on air pollutant-induced cardiopulmonary and vascular pathologies, next we investigated the effects of haplo- and complete genetic deficiency of PAI-1 on air pollutant PM_2.5_-induced cardiopulmonary pathologies. Wild-type, PAI-1 heterozygous, and PAI-1 knockout mice were treated with PM_2.5_ (50 μg/mouse/week for 4 weeks) or PBS as a control. All mice were fed with regular chow. Four weeks of PM_2.5_ exposure induces pulmonary thrombosis in both wild-type and PAI-1 knockout mice, indicating complete deficiency of PAI-1 does not protect against developing PM_2.5_-induced pulmonary pathologies. Interestingly, PAI-1 heterozygous mice are partially protected from PM_2.5_-induced pulmonary thrombotic pathogenesis, indicating that, like pharmacological inhibition of PAI-1, PAI-1 haplodeficiency, but not complete absence of PAI-1, is beneficial for air pollutant-induced pulmonary pathologies (Supplemental Figure s7). However, we did not observe any difference in pulmonary matrix remodeling in PM_2.5_-treated wild-type, PAI-1 heterozygous, and knockout groups compared to controls and TM5614-treated groups, as evidenced by Masson’s trichrome staining (*data not shown*).

### TM5614 reduces PM_2.5_-induced cellular stresses and abnormalities

As PM_2.5_ induces the levels of reactive oxygen species, TGF-β, and matrix remodeling, and fibroblasts are the major contributing cell type in matrix protein production, we have developed an *in vitro* model system to study the effects of PM_2.5_ on cellular processes in cardiac fibroblasts and the efficacy of TM5614 in ameliorating PM_2.5_-induced cellular abnormalities. Our results showed that PM_2.5_ modestly induces the levels of PAI-1, type I collagen, and fibronectin and that TM5614 reduces the PM_2.5_-induced stimulation of these fibrogenic factors (Figure 10A-C). The PM_2.5_-induced stimulation of PAI-1 and its normalization by TM5614 *in vitro* is consistent with the *in vivo* observation on plasma levels of PAI-1 in PM_2.5_- and TM5614-treated mice (Figure 1). As SREBP1, a basic-helix-loop-helix leucine zipper family transcription factor, mediates profibrogenic signaling and fibrogenesis,^24^ and air pollutants induce the expression of SREBP1,^25^ we measured the levels of SREBP1 and SREBP2 in control, PM_2.5_-, and TM5614-treated cardiac fibroblasts. Interestingly, PM_2.5_ induces the expression of SREBP1/2 in human cardiac fibroblasts and most importantly, TM5614 completely abrogates that stimulation, indicating that TM5614 effectively blocks PM_2.5_-induced profibrogenic signaling through suppression of SREBP1 and SREBP2 (Figure 10A). As PM_2.5_ induces reactive oxygen species^26^ and Nrf2, a basic leucine zipper (bZip) transcription factor, is a master regulator of antioxidant genes and signaling,^27^ we measured the effect of PM_2.5_ on Nrf2 expression in human cardiac fibroblasts and the efficacy of TM5614 in normalizing the Nrf2 expression. Our results revealed that PM_2.5_ reduces the levels of Nrf2 in cardiac fibroblasts and importantly, that TM5614 reverses the suppression of Nrf2 by PM_2.5_ (Figure 10C). Collectively, these *in vitro* results indicate the efficacy of PAI-1 inhibitor TM5614 in improvement of cardiac cellular stresses and abnormalities induced by exposure to air pollutant PM_2.5_.

**Figure 10.**
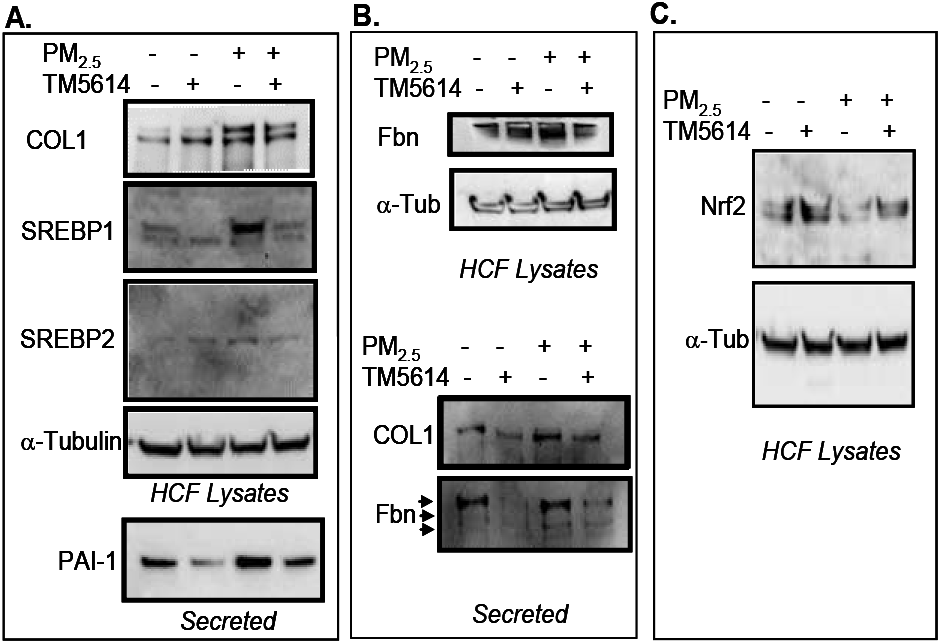
TM5614 blocks PM_2.5_-induced cellular stress and profibrogenic responses. Human cardiac fibroblasts were cultured in 12-well clusters. After 24 h, media, were replaced with 0.1% FBS containing DMEM for 3 h. Cells were pretreated with TM5614 (10 μM) or (vehicle control) for 2 h followed by treatment with PM_2.5_ (50 μg/ml) in triplicate for another 2 h. Supernatants and cell lysates were used for western blotting using antibodies as indicated (**A**). In (**B**) and (**C**), cells were pretreated with TM5614 (10 μM) or in triplicate. After 24 h, media with replaced with 0.1% FBS containing and treated with TM5614 (10 μM) or and PM_2.5_ (50 μg/ml) in triplicate for another 24 h. Supernatants and cell lysates were processed for western blotting using antibodies as indicated (**B,C**).

## DISCUSSION

Environmental pollutants like PM are a global threat to organismal growth and health. Short-term or long-term exposure to environmental pollutants like PM causes devastating CPVDs. CPVDs including pulmonary inflammation, vascular dysfunction, hypertension, arteriosclerosis, arrhythmias, and myocardial infarction cause an estimated 60-80% of environmental pollutant-associated deaths worldwide.^1–8^ However, the underlying molecular basis of PM-induced CPVDs is not well understood, and an effective therapeutic approach to block PM-induced CPVD-related pathologies is not available. As exposure to environmental pollutants deregulates the plasminogen activator system,^15,16^ and PAI-1 plays pivotal roles in initiation and progression of CPVDs,^10–14^ we initiated the present study in order to delineate the role of PAI-1 in air pollutant PM_2.5_-induced CPVD-related pathologies, and assess the potential of PAI-1 as a druggable target for CPVD therapy. Using the small molecule inhibitor TM5614, which targets PAI-1, and mouse strains with PAI-1 haplo- or complete deficiency, the present study demonstrated that PAI-1 plays a key role in air pollutant PM_2.5_-induced CPVD pathologies and that pharmacological inhibition of PAI-1 ameliorates PM_2.5_-induced CPVD pathologies.

In this study, we demonstrated that short-term exposure to air pollutant PM_2.5_ increases the levels of pro-inflammatory and pro-thrombotic PAI-1 and thrombin in plasma and inflammatory cytokine IL-6 in lungs, consistent with previous observations.^15^ The levels of TAT complex serve as an indicator of coagulation activation. Thrombin is a serine protease that converts soluble fibrinogen to an insoluble fibrous protein, fibrin, that polymerizes and becomes a key component of blood clots. Antithrombin, an endogenous inhibitor of thrombin, forms the TAT complex with thrombin. As thrombin is highly unstable and has a short half-life in plasma, the TAT complex is measured as an indicator of thrombin levels and activation of coagulation that contributes to thrombosis.^28^

Most importantly, here we have presented data showing that the stimulation of PAI-1, thrombin, and IL-6 in air pollutant-exposed mice is abrogated by treatment with PAI-1 inhibitor TM5614. The elevated levels of PAI-1 and IL-6 clearly indicate increased inflammation by exposure to air pollutant PM_2.5_, and early inflammation plays a pivotal role in the development of downstream inflammatory signals and organ pathologies, especially in the lungs and cardiovascular system, including vascular occlusion and disruption of lung and heart tissue homeostasis (Figure 11). Suppression of IL-6 by PAI-1 inhibitor TM5614 can be explained in the light of previous observations made by Kubala and colleagues,^29^ who demonstrated that while elevated PAI-1 stimulates the levels of IL-6 and activates downstream pSTAT3, depletion of PAI-1 is associated with down-regulation of IL-6 and pSTAT3 activation in response to inflammatory signals.^29^

**Figure 11.**
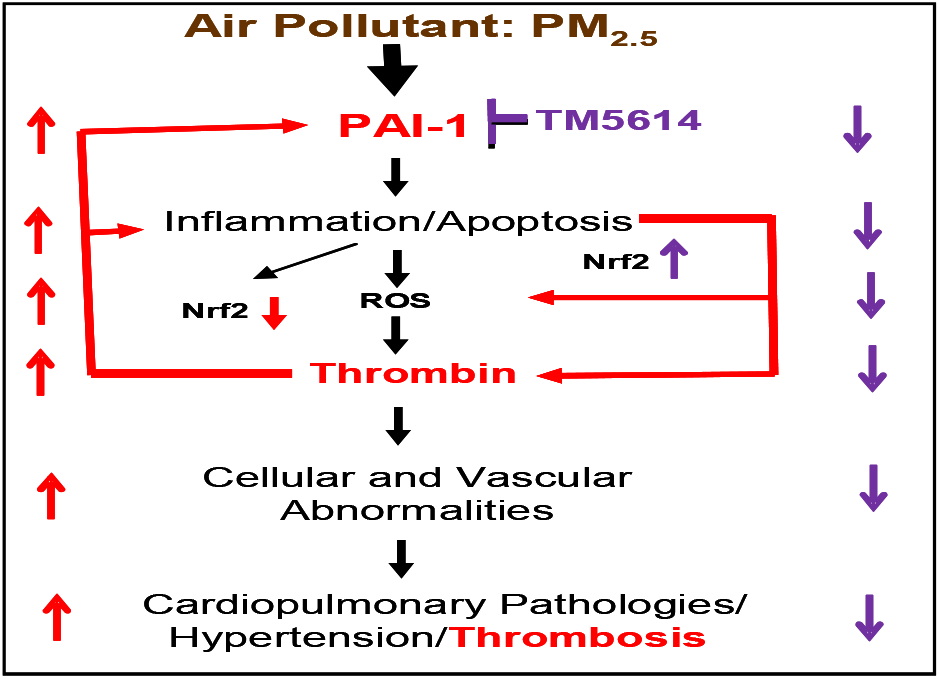
Possible pathological pathways driven by PM_2.5_-induced PAI-1 and the beneficial effect of PAI-1 inhibitor TM5614. Air pollution stressors increase the levels of proinflammatory and prothrombotic mediators/ regulators that cause cellular and vascular dysfunction and contribute to cardiopulmonary vascular pathologies. Neutralization of PM_2.5_-induced PAI-1 with TM5614 reduces PM_2.5_-induced cardiopulmonary vascular pathologies.

Next, we investigated the effect of short-term exposure to PM_2.5_ on lung and heart inflammation and apoptosis and showed that the lung morphology is abnormal in several PM_2.5_-exposed mice (200 μg/mouse) in terms of its tissue density, consistent with previous observations that PM_2.5_ causes damage in lung morphological structure.^30^ These results indicate that acute exposure to air pollutant is able to damage lung tissue architecture. However, in the presence of PAI-1 inhibitor TM5614, PM_2.5_ fails to induce pulmonary tissue damage. Furthermore, TM5614 reduces air pollutant PM_2.5_-induced inflammation, apoptosis, and vascular dysfunction as evidenced by the decreased levels of inflammation- and apoptosis-associated factors in lungs derived from mice exposed to PM_2.5_ in the presence of TM5614. Similarly, TM5614 is also effective in suppression of PM_2.5_-induced modestly elevated inflammatory and apoptotic events in murine hearts. These results signify that pharmacological neutralization of PM_2.5_-induced PAI-1 is an effective approach for suppression of early events of air pollutant-induced cardiovascular and pulmonary pathological events. Previous studies demonstrated that PM_2.5_ induces the levels of PAI-1^15,16^ and that elevated PAI-1 plays a key role in a number of subclinical and clinical pathologies, including inflammation, atherosclerosis, insulin resistance, obesity, and multimorbidities.^10,11,14,31^ The elevated PAI-1 may also contribute to cellular apoptosis in a context-dependent manner. For example, the elevated PAI-1 in alveolar type II cells derived from idiopathic pulmonary fibrosis patients is associated with elevated levels of cleaved caspase 3 and thus increased apoptosis.^32^ Therefore, pharmacological suppression or neutralization of PAI-1 is an ideal approach to control pathological/sustained inflammation and disease development like air pollutant-induced increased inflammation and apoptosis-driven cardiopulmonary diseases.

The most notable observation we made in the present study is the induction of significant levels of pulmonary and, to a lesser degree, cardiac vascular thrombosis in mice exposed to air pollutant PM_2.5_, and that air pollutant exposure fails to develop pulmonary and cardiac thrombosis in the mice treated with PAI-1 inhibitor TM5614. This result is consistent with our observation that TM5614 significantly reduces PM_2.5_-induced increased levels of inflammation and pro-thrombotic mediators IL-6, PAI-1, and thrombin. Furthermore, normalization of PM_2.5_-induced increased blood pressure and heart weight by TM5614 treatment are also significant in the context of the pivotal role of PAI-1 in PM_2.5_-induced vascular pathologies. In this context, it is important to note that previous studies implicated PM and higher levels of PAI-1 in high blood pressure.^33,34^ Like wild-type mice, several PAI-1 knockout mice also developed pulmonary thrombosis upon longer exposure to air pollutant PM_2.5_. Interestingly, PAI-1 haplodeficient mice exposed to PM_2.5_ are partly protected from developing pulmonary thrombosis, indicating that, like pharmacological inhibition of PAI-1, partial genetic deficiency of PAI-1 or low levels of PAI-1 are beneficial for organisms exposed to air pollutants. Therefore, it is apparent that normalization of the levels of PAI-1, but not complete depletion, is favorable to prevent air pollutant-induced pulmonary pathologies. Further studies are necessary to understand the underlying molecular basis. Although previous reports suggested significant cardiac matrix remodeling in mice and rats in response to exposure to air pollution,^35–37^ under our experimental setup (100 μg/once/week/4 weeks), we did not observe any significant change in collagen deposition in PM_2.5_-exposed murine lungs and hearts. However, our *in vitro* study showed up-regulation of profibrogenic signal transducers and matrix proteins in cardiac fibroblasts exposed to PM_2.5_, and importantly, TM5614 is effective in the suppression of these PM_2.5_-induced profibrogenic responses. Further studies are required to investigate the effect of PM_2.5_ exposure with an increased number of doses/week for 3-6 months on pulmonary and cardiac pathological matrix remodeling and whether TM5614 can effectively block PM_2.5_-induced pulmonary and cardiac fibrogenesis on that timescale. Together, these results signify the pivotal role of PAI-1 in air pollutant-induced cardiopulmonary vascular pathologies and establish the potential therapeutic efficacy of PAI-1 inhibitor TM5614, an FDA-approved drug for clinical trials of other PAI-1 related pathologies, in amelioration of air pollutant exposure-mediated cardiopulmonary pathologies.

In search of cellular and molecular events induced by exposure to air pollutant PM_2.5_, we studied the effects of PM_2.5_ on cultured human cardiac fibroblasts in the presence and absence of TM5614. The *in vitro* results suggest that air pollutant PM_2.5_ modestly induces profibrogenic markers and regulators, namely profibrogenic PAI-1, matrix protein collagen, and transcriptional regulators SREBP1 and SREBP2; and most importantly, that PAI-1 inhibitor efficiently blocks air pollutant-induced increases in PAI-1, profibrogenic regulators, and target matrix proteins. The rescue of PM_2.5_-induced suppression of Nrf2 levels by TM5614 is interesting, as it clearly indicates that suppression of Nrf2, a major transcriptional regulator of antioxidant genes, and augmentation of oxidative stress by PM_2.5_ can be prevented by pharmacological inhibition of PAI-1. This result is consistent with our previous observations showing PAI-1 inhibition is associated with the rescue of homocysteine-mediated suppression of the antioxidant regulator Nrf2^13^ and of doxorubicin-mediated suppression of antioxidant catalase,^12^ as well as with prevention of accelerated cellular aging.

In conclusion, PAI-1 is a potentially druggable target for the treatment of air pollutant-induced cardiopulmonary vascular pathologies including inflammation, thrombosis, hypertension, and cardiac hypertrophy; and TM5614, a drug-like small molecule inhibitor of PAI-1, effectively ameliorates these cardiopulmonary vascular pathologies by suppression of air pollutant PM_2.5_-induced regulators of inflammation, prothrombotic factors, profibrogenic regulators, and by rescue of PM_2.5_-induced suppression of a key antioxidant regulator (Figure 11). Further *in vitro* studies are needed to test the efficacy of TM5614 in amelioration of other air pollutant-induced signaling pathways involved in cardiopulmonary vascular pathologies, including inflammatory signaling, canonical or non-canonical TGF-β signaling, and pro-apoptotic signaling pathways in endothelial cells and macrophages. Future expansion of the present study will be helpful to design clinical trials of TM5614 for the treatment of air pollutant-induced CPVDs.

## Acknowledgments

We thank Northwestern University’s Mouse Histology and Phenotyping Laboratory for technical support.

## Source of Funding

This work is supported by the American Heart Association-Innovative Project Award (18IPA34170365) and by a grant from the National Institutes of Health (5R01HL051387).

## Disclosures

None

## Ghosh et al. Supplemental Data

**Supplemental Figure s1.**
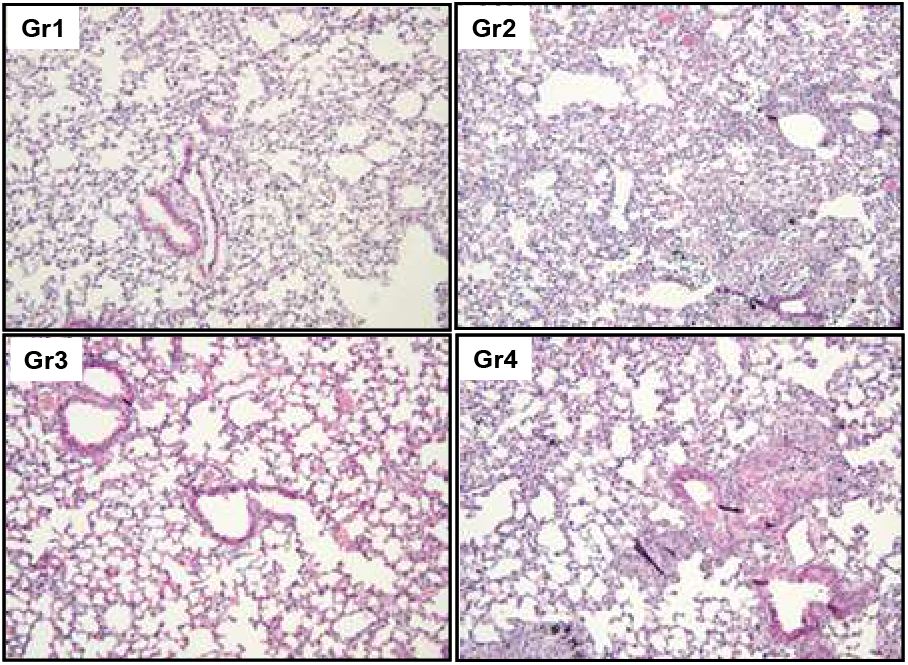
Effects of PM_2.5_ and PAI-1 inhibitor TM5614 on lung morphology. Lungs collected from 4-groups were processed for H&E staining (n=4-7). Day 1-6: TM5614 (10 mg/Kg) or normal diet; Day 7: PM_2.5_ (200 μg/50 μl/mouse) or PBS instillation; Day 10: Lungs collected were processed for morphology study. Images reduced to 15% of original 20X images. Gr1 (PBS); Gr2: (PM_2.5_); Gr3: TM5614; Gr4: PM_2.5_+TM5614. Morphology denser: Gr1: 1/4; Gr2: 4/7; Gr3: 1/4; Gr4: 2/6

**Supplemental Figure s2.**
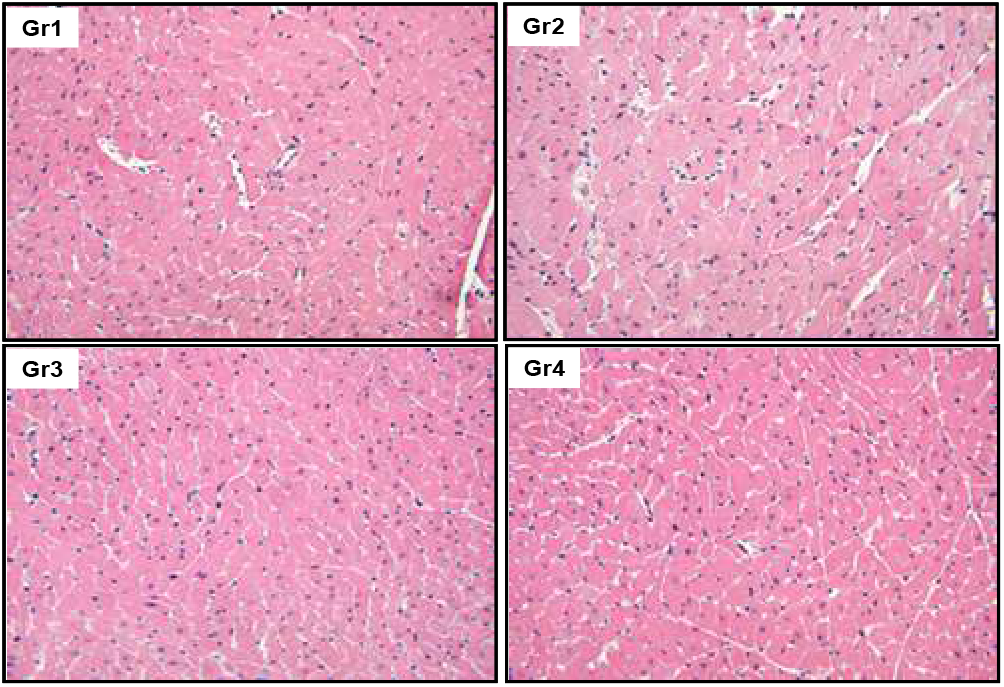
Effect of PM_2.5_ and PAI-1 inhibitor TM5614 on cardiac morphology. Hearts collected from 4-groups (n=4-7) as indicated were processed for haematoxylin and eosin staining. [Day 1-6: TM5614 (10 mg/Kg); Day 7: PM_2.5_ (200 μg/mouse) or PBS instillation. Day 10: Hearts collected and processed for histological analysis. Images reduced to 15% of original 40X magnification. Gr1: PBS; Gr2: PM_2.5_; Gr3: TM5614; Gr4: PM_2.5_+TM5614.

**Supplemental Figure s3.**
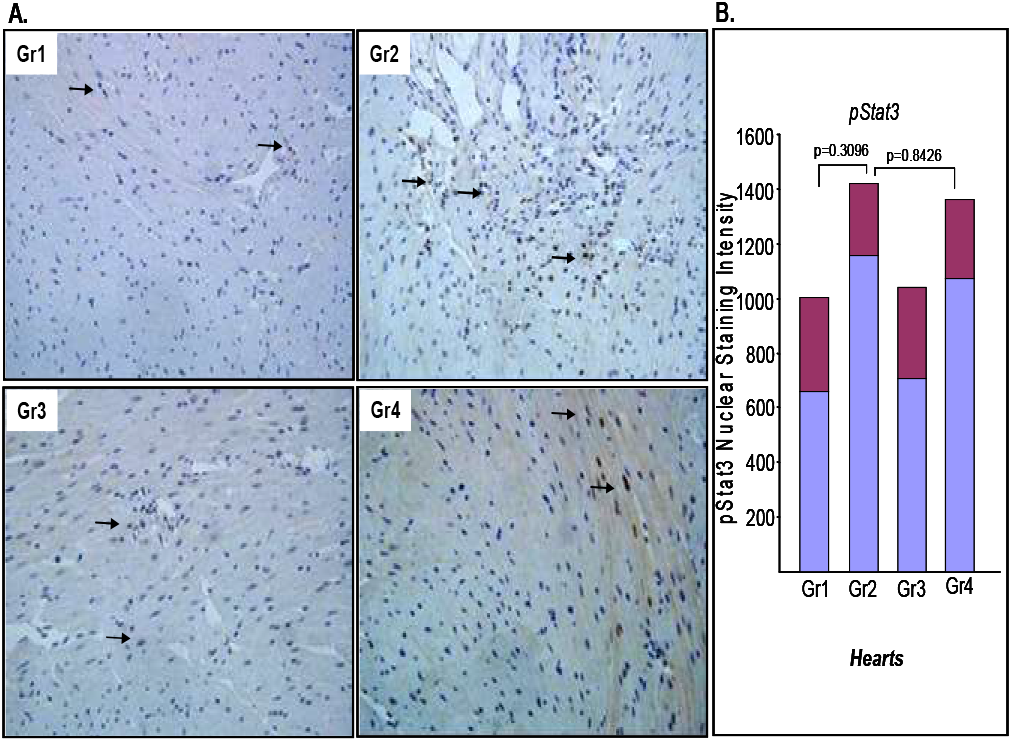
Effect of PM_2.5_ and PAI-1 inhibitor TM5614 on inflammatory factor pStat3 in Hearts. Hearts were collected from 4-groups (n=3-6) and formalin fixed. Heart tissues were processed for immunohistochemistry using inflammatory marker pSTAT3 antibody. [Day 1-6: TM5614 (10 mg/Kg); Day 7: PM_2.5_ (200 μg/50 ml/mouse/once) or PBS intubation. Day 10: Hearts were collected and processed for immunohistochemistry. The representative image from each group was presented in (**A**). The levels of nuclear pSTAT3 in several fields of heart sections were determined by ImageJ software followed by statistical analysis (**B**). Representative images reduced to 15% of original 40X magnification. Gr1(PBS); Gr2 (PM_2.5_); Gr3 (TM5614); Gr4 (TM5614+PM_2 5_).

**Supplemental Figure s4.**
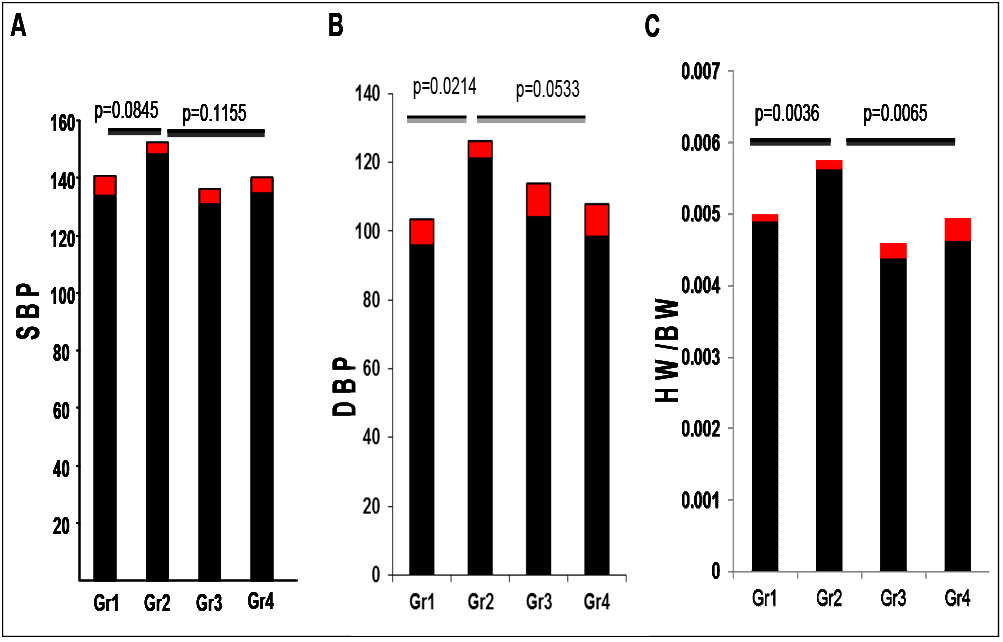
Effect of long-term exposure to PM_2.5_-on blood pressure and cardiac hypertrophy: Mice were divided into 4 groups. In Group1, mice were fed with regular chow and received PBS. In Group 2, mice were fed with regular chow and received PM_2.5_ (100 μg/mouse/once per week for 4 weeks). In Group 3, mice were fed with chow containing TM5614 (10 mg/kg) for 6 days and then received PBS. In Group 4, mice were fed with chow containing TM5614 (10 mg/kg) for 6 days and then received PM_2.5_ (100 μg/mouse/once per week for 4 weeks). Systolic (**A**) (n=3-7) and diastolic (**B**) (n=3-7) blood pressure in four groups. Heart weight/body weight in four groups are shown in (**C**). n=4-7.

**Supplemental Figure s5.**
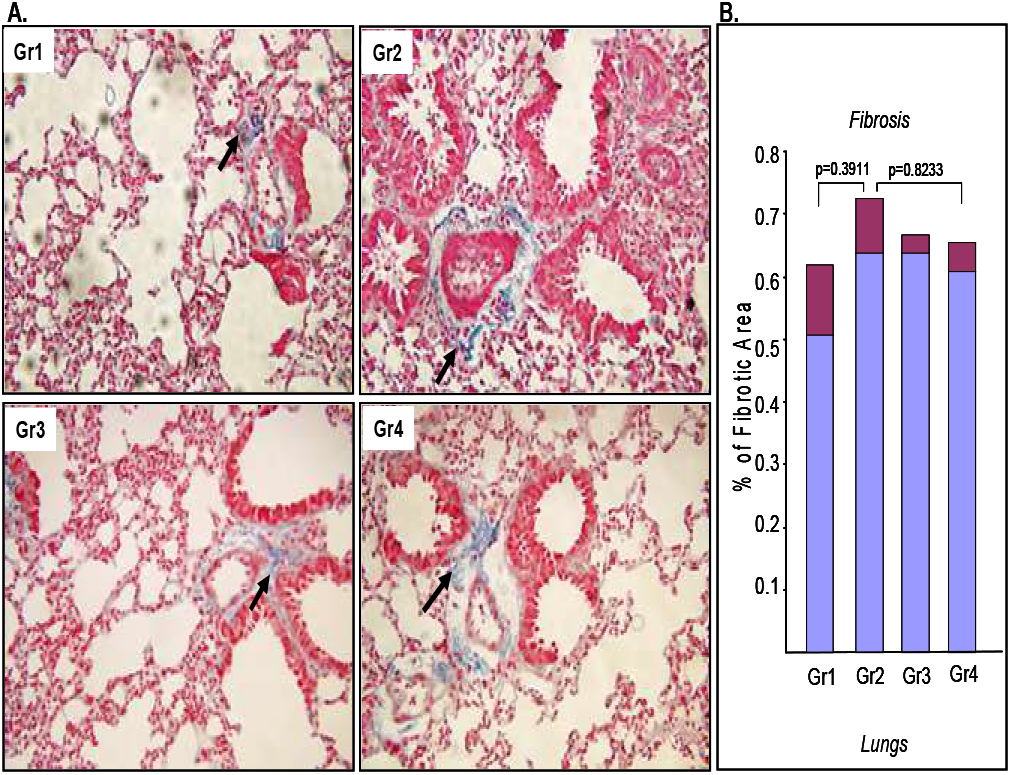
Effects of PM_2.5_ and PAI-1 inhibitor TM5614 on pulmonary matrix remodeling. Lungs collected from 4-groups (n=4-7) were processed for Masson’s trichrome staining to determine the levels of pulmonary fibrosis. Mice were divided into 4 groups. In Group1, mice were fed with regular chow and received PBS as control. In Group 2, mice were fed with regular chow and received PM_2.5_ (100 μg/mouse/once per week for 4 weeks). In Group 3, mice were fed with chow containing TM5614 (10 mg/kg) for 6 days and then received PBS. In Group 4, mice were fed with chow containing TM5614 (10 mg/kg) for 6 days and then received PM_2.5_ (100 μg/mouse/once per week for 4 weeks). Lungs collected were processed for Masson’s trichrome staining. Representative images are reduced to 15% of original 40X images (A). Quantitative data of fibrosis using Fiji is just ImageJ software are shown in B.

**Supplemental Figure s6.**
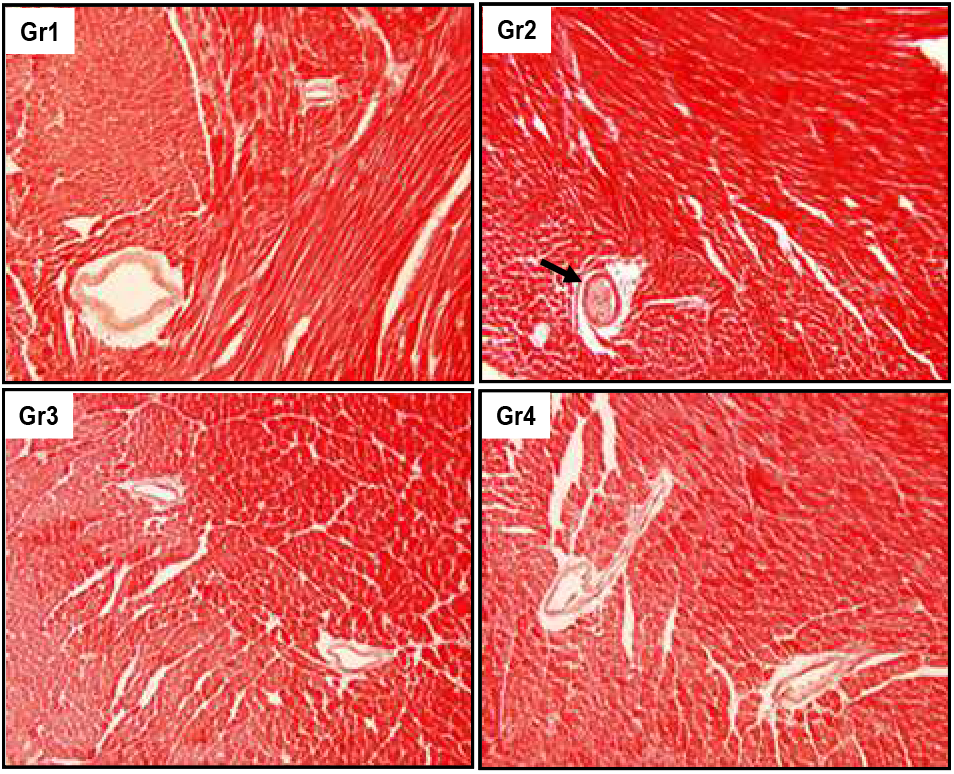
Effects of TM5614 on PM_2.5_-induced cardiac thrombosis. Mice were divided into 4 groups. In Group1, mice were fed with regular chow and received PBS. In Group 2, mice were fed with regular chow and received PM_2.5_ (100 μg/mouse/once per week for 4 weeks). In Group 3, mice were fed with chow containing TM5614 (10 mg/kg) for 6 days and then received PBS. In Group 4, mice were fed with chow containing TM5614 (10 mg/kg) for 6 days and then received PM_2.5_ (100 μg/mouse/once per weeks for 4 weeks). Analysis of hearts from each group reveal that exposure to PM_2.5_ modestly induces cardiac microthrombi with lesser extent compared to lungs in mice (microthrombi in Gr1: 0/4 mice; Gr2: 4/7 mice; Gr3: 0/5 mice; Gr4:0/4 mice). PM_2.5_ fails to induce cardiac thrombosis in mice treated with PAI-1 inhibitor TM5614 (Gr4). Images reduced to 15% of original 20X images are shown.

**Supplemental Figure s7.**
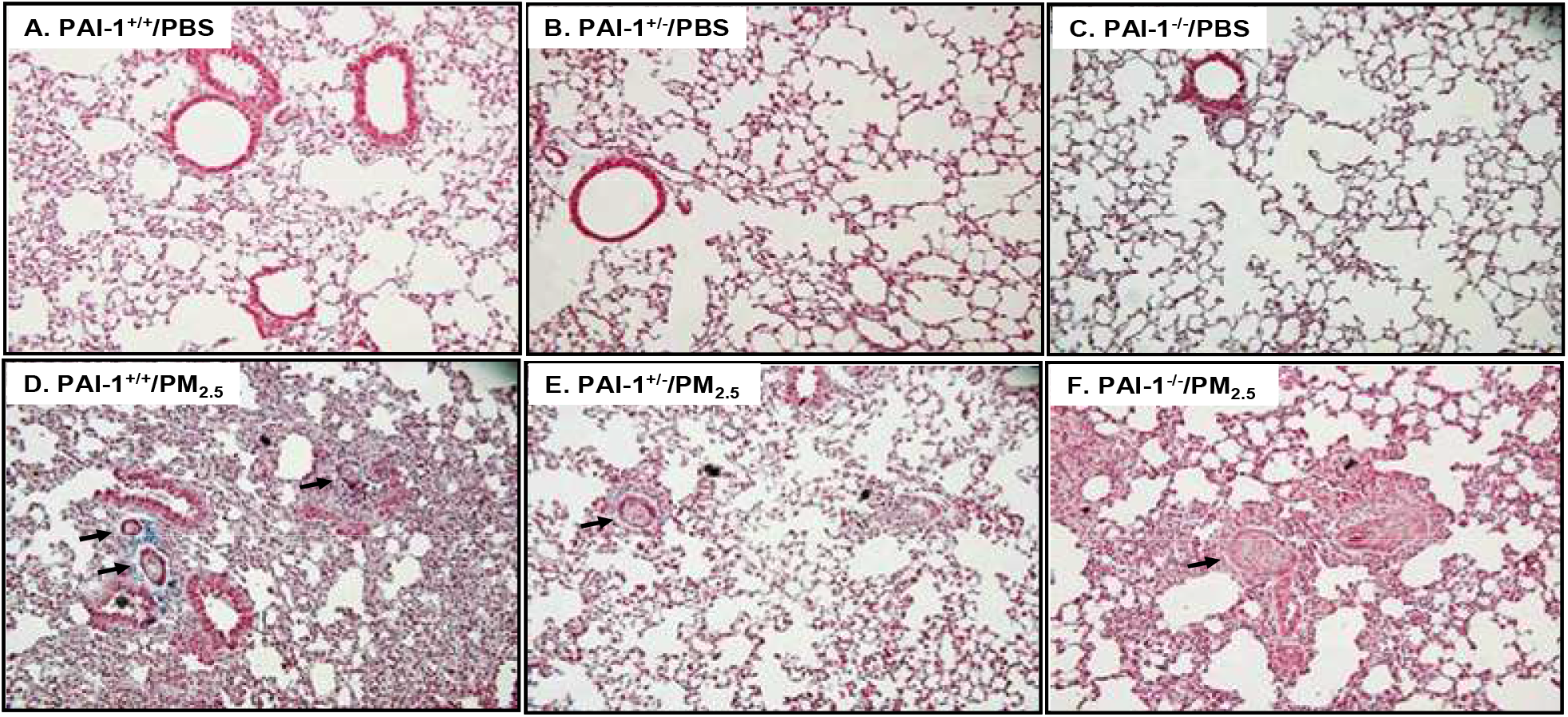
Effects of PAI-1 haplo and complete deficiency on air pollutant PM_2.5_-induced pulmonary pathologies. The wildtype, PAI-1 heterozygous and PAI-1 knockout mice were treated with PM_2.5_ (50μg/mouse/week for 4 weeks) or PBS as controls. All mice were fed with regular chow. Each group of mice viz. wildtype (A,D), PAI-1 heterozygous (B,E) and PAI-1 global knockout mice (C,F) were divided into 2 groups. While Group1 mice received 50 μl of PBS (A, B, C), Group 2 mice received PM_2.5_ (50 μg/mouse/once per week for 4 weeks) (D, E, F). There are total 6 groups. Representative images showing microthrombi are presented in A. Images reduced to 15% of original 20X magnification. Arrows indicate PM_2.5_-induced microthrombi in lungs. Microthrombi in A:0/5; B:0/6; C: 0/4; D: 4/5; E:1/4; F: 3/6 mice.

## Notes

### Competing Interest Statement

The authors have declared no competing interest.

## REFERENCES

1. Bourdrel T, Bind MA, Béjot Y, Morel O, Argacha JF. Cardiovascular effects of air pollution. Arch Cardiovasc Dis.2017;110:634–642.

2. Du Y, Xu X, Chu M, Guo Y, Wang J. Air particulate matter and cardiovascular disease: the epidemiological, biomedical and clinical evidence. J Thorac Dis. 2016;8:E8–E19.

3. Brook RD, Franklin B, Cascio W, Hong Y, Howard G, Lipsett M, Luepker R, Mittleman M, Samet J, Smith SC Jr, Tager I. Expert Panel on Population and Prevention Science of the American Heart Association. Air pollution and cardiovascular disease: a statement for healthcare professionals from the Expert Panel on Population and Prevention Science of the American Heart Association. Circulation. 2004;109:2655–2671.

4. Brook RD, Rajagopalan S, Pope CA 3rd, Brook JR, Bhatnagar A, Diez-Roux AV, Holguin F, Hong Y, Luepker RV, Mittleman MA, Peters A, Siscovick D, Smith SC Jr, Whitsel L, Kaufman JD. American Heart Association Council on Epidemiology and Prevention, Council on the Kidney in Cardiovascular Disease, and Council on Nutrition, Physical Activity and Metabolism. Particulate matter air pollution and cardio-vascular disease: An update to the scientific statement from the American Heart Association. Circulation. 2010;121:2331–2378.

5. Nel A. Atmosphere. Air pollution-related illness: effects of particles. Science. 2005;308:804–806.

6. Franchini M, Mannucci PM. Thrombogenicity and cardiovascular effects of ambient air pollution. Blood. 2011;118:2405–2412.

7. Budinger GR, Mutlu GM. Update in environmental and occupational medicine 2010. Am J Respir Crit Care Med. 2011;183:1614–1619.

8. Simkhovich BZ, Kleinman MT, Kloner RA. Particulate air pollution and coronary heart disease. Curr Opin Cardiol. 2009;24:604–609.

9. Robertson S, Miller MR. Ambient air pollution and thrombosis. Part Fibre Toxicol. 2018;3;15:1.

10. Westrick RJ, Eitzman DT. Plasminogen activator inhibitor-1 in vascular thrombosis. Curr Drug Targets. 2007;8:966–1002.

11. Ghosh AK, Vaughan DE. PAI-1 in Tissue Fibrosis. J Cell Physiol. 2012; 227: 493–507.

12. Ghosh AK, Rai R, Park KE, Eren M, Miyata T, Wilsbacher LD, Vaughan DE. A small molecule inhibitor of PAI-1 protects against doxorubicin-induced cellular senescence. Oncotarget. 2016;7:72443–72457.

13. Sun T, Ghosh AK., Eren M, Miyata T, Vaughan DE. PAI-1 contributes to Homocysteine-induced Cellular Senescence. Cell Signal. 2019;64:109394.

14. Vaughan DE, Rai R, Khan SS, Eren M, Ghosh AK. Plasminogen Activator Inhibitor-1 Is a Marker and a Mediator of Senescence. Arterioscler Thromb Vasc Biol. 2017;37:1446–1452.

15. Budinger GR, McKell JL, Urich D, Foiles N, Weiss I, Chiarella SE, Gonzalez A, Soberanes S, Ghio AJ, Nigdelioglu R, Mutlu EA, Radigan KA, Green D, Kwaan HC, Mutlu GM. Particulate matter-induced lung inflammation increases systemic levels of PAI-1 and activates coagulation through distinct mechanisms. PLoS One. 2011;6:e18525.

16. Upadhyay S, Ganguly K, Stoeger T, Semmler-Bhenke M, Takenaka S, Kreyling WG, Pitz M, Reitmeir P, Peters A, Eickelberg O, Wichmann HE, Schulz H. Cardiovascular and inflammatory effects of intratracheally instilled ambient dust from Augsburg, Germany, in spontaneously hypertensive rats (SHRs). Part Fibre Toxicol. 2010;7:27.

17. Soberanes S, Urich D, Baker CM, Burgess Z, Chiarella SE, Bell EL, Ghio AJ, De Vizcaya-Ruiz A, Liu J, Ridge KM, Kamp DW, Chandel NS, Schumacker PT, Mutlu GM, Budinger GR. Mitochondrial complex III-generated oxidants activate ASK1 and JNK to induce alveolar epithelial cell death following exposure to particulate matter air pollution. J Biol Chem. 2009;284:2176–2186.

18. Yang J, Chen Y, Yu Z, Ding H, Ma Z. The influence of PM_2.5_ on lung injury and cytokines in mice. Exp Ther Med 2019;18:2503–2511.

19. Pei Y, Jiang R, Zou Y, Wang Y, Zhang S, Wang G, Zhao J, Song W. Effects of Fine Particulate Matter (PM_2.5_) on Systemic Oxidative Stress and Cardiac Function in ApoE^-/-^ Mice. Int J Environ Res Public Health. 2016, 13, 484.

20. Huang K, Shi C, Min J, Li L, Zhu T, Yu H, Deng H. Study on the Mechanism of Curcumin Regulating Lung Injury Induced by Outdoor Fine Particulate Matter (PM_2.5_). Mediators Inflamm. 2019;2019:8613523.

21. Jeong S, Park SA, Park I, Kim P, Cho N-H, Hyun JW, Hyun Y-M. PM2.5 Exposure in the Respiratory System Induces Distinct Inflammatory Signaling in the Lung and the Liver of Mice. J Immunol Res, 2019, 11

22. Cui A, Xiang M, Xu M, Lu P, Wang S, Zou Y, Qiao K, Jin C, Li Y, Lu M, Chen AF, Chen S. VCAM-1-mediated neutrophil infiltration exacerbates ambient fine particle-induced lung injury. Toxicol Lett. 2019;302:60–74

23. Rui W, Guan L, Zhang F, Zhang W, Ding W. PM2.5-induced oxidative stress increases adhesion molecules expression in human endothelial cells through the ERK/AKT/NF-κB dependentpathway. J Appl Toxicol. 2016;36:48–59.

24. Chen G, Wang T, Uttarwar L, vanKrieken R, Li R, Chen X, Gao B, Ghayur A, Margetts P, Krepinsky JC. SREBP-1 is a novel mediator of TGFβ1 signaling in mesangial cells. J Mol Cell Biol. 2014;6:516–30.

25. Yan R, Ku T, Yue H, G Nan, Sang N. PM2.5 exposure induces age-dependent hepatic lipid metabolism disorder in female mice. J Environ Sci 2020; 89: 227–237.

26. Liu CW, Lee TL, Chen YC, Liang C-J, Wang S-H, Lue J-H, Tsai J-S, Lee S-W, Chen S-H, Yang Y-F, Chuang T-Y, Chen Y-L. PM_2.5_-induced oxidative stress increases intercellular adhesion molecule-1 expression in lung epithelial cells through the IL-6/AKT/STAT3/NF-κB-dependent pathway. Part Fibre Toxicol 2018;15:4.

27. Vomund S, Schäfer A, Parnham MJ, Brüne B, von Knethen A. Nrf2, the Master Regulator of Anti-Oxidative Responses. Int J Mol Sci. 2017; 18:2772.

28. Chen Q, Shou W, Wu W, Wang G, Cui W Performance evaluation of thrombomodulin, thrombin-antithrombin complex plasmin-α2-antiplasmin complex, and t-PA: PAI-1 complex. .J Clin Lab Anal. 2019;33:e22913.

29. Kubala MH, Punj V, Placencio-Hickok VR, Fang H, Fernandez GE, Sposto R, DeClerck YA Plasminogen Activator Inhibitor-1 Promotes the Recruitment and Polarization of Macrophages in Cancer. Cell Rep. 2018;25:2177–2191.

30. Yang J, Chen Y, Yu Z, Ding H, Ma Z. The influence of PM_2.5_ on lung injury and cytokines in mice. Exp Ther Med. 2019;18:2503–2511.

31. Cesari M, Pahor M, Incalzi RA. Plasminogen activator inhibitor-1 (PAI-1): a key factor linking fibrinolysis and age-related subclinical and clinical conditions. Cardiovasc Ther. 2010;28:e72–91.

32. Marudamuthu AS, Shetty SK, Bhandary YP, Karandashova S, Thompson M, Sathish V, Florova G, Hogan TB, Pabelick CM, Prakash YS, Tsukasaki Y, Fu J, Ikebe M, Idell S, Shetty S Plasminogen activator inhibitor-1 suppresses profibrotic responses in fibroblasts from fibrotic lungs. J Biol Chem. 2015; 290:9428–41.

33. Peng H, Yeh F, de Simone G, Best LG, Lee ET, Howard BV, Zhao J. Relationship Between Plasma Plasminogen Activator inhibitor-1 and Hypertension in American Indians: Findings From the Strong Heart Study. J Hypertens. 2017; 35:1787–1793.

34. Brook RD, Rajagopalan S. Particulate matter, air pollution, and blood pressure. J Am Soc Hypertens. 2009; 3:332–350.

35. Xu Z, Li, Z, Liao Z, Gao S, Hua L, Ye X, Wang Y, Jiang S, Wang N, Zhou D, Deng X. PM_2.5_Induced Pulmonary Fibrosis in Vivo and in Vitro. Ecotoxicol Environ Saf 2019;171:112–121.

36. Sun B, Shi Y, Li Y, Jiang J-j, Liang S, Duan J, Sun Z. Short-term PM2.5 exposure induces sustained pulmonary fibrosis development during post-exposure period in rats. J Hazard Mater, 2020; 385:121566.

37. Yue W, Tong L, Liu X, Weng X, Chen X, Wang D, Dudley SC, Weir EK, Ding W, Lu Z, Xu Y, Chen Y. Short term PM2.5 exposure caused a robust lung inflammation, vascular remodeling, and exacerbated transition from left ventricular failure to right ventricular hypertrophy. Redox Biol. 2019;22:101161.

